# AMPK signaling regulates expression of urea cycle enzymes in response to changes in dietary protein intake

**DOI:** 10.1101/439380

**Authors:** Sandra Kirsch Heibel, Peter J McGuire, Nantaporn Haskins, Himani Datta Majumdar, Sree Rayavarapu, Kanneboyina Nagaraju, Yetrib Hathout, Kristy Brown, Mendel Tuchman, Ljubica Caldovic

## Abstract

Abundance of urea cycle enzymes in the liver is regulated by the dietary protein intake. Although urea cycle enzyme levels rise in response to a high protein diet, signaling networks that sense dietary protein intake and trigger changes in expression of urea cycle genes have not been identified. The aim of this study was to identify signaling pathway(s) that respond to changes in protein intake and regulate expression of urea cycle genes in mice and human hepatocytes. Mice were adapted to either control or high (HP) protein diets followed by isolation of liver protein and mRNA and integrated analysis of the proteomic and transcriptome profiles. HP diet led to increased expression of mRNA and enzymes in amino acid degradation pathways, and decreased expression of mRNA and enzymes in carbohydrate and fat metabolism, which implicated AMPK as a possible regulator. Primary human hepatocytes, treated with AICAR an activator of AMPK, were used to test whether AMPK regulates expression of urea cycle genes. The abundance of *CPS1* and *OTC* mRNA increased in hepatocytes treated with AICAR, which supports a role for AMPK signaling in regulation of the urea cycle. Because AMPK is either a target of drugs used to treat type-2 diabetes, these drugs might increase the expression of urea cycle enzymes in patients with urea cycle disorders, which could be the basis of a new therapeutic approach.

**Author summary:** Integrated analysis of transcriptional and proteomic profiles of the liver tissue from mice fed different protein content diets revealed that AMPK signaling pathway regulates expression of urea cycle enzymes.

## Introduction

Six enzymes, N-acetylglutamate synthase (NAGS; EC 2.3.1.1), carbamylphosphate synthetase 1 (CPS1, EC 6.3.4.16), ornithine transcarbamylase (OTC, EC 2.1.3.3), argininosuccinate synthase (ASS, EC 6.4.3.5), argininosuccinate lyase (ASL, EC 4.3.2.1), and arginase 1 (ARG1, EC 3.5.3.1), and two transporters, ornithine/citrulline transporter (ORNT) and aspartate/glutamate transporter (ARALAR2 or Citrin) comprise the urea cycle in the liver [1, 2]. The physiological function of the urea cycle is to convert ammonia, a neuro-toxic product of protein catabolism, into urea. Defects in any of the urea cycle enzymes and transporters, collectively known as urea cycle disorders (UCD), lead to hyperammonemia, which causes neuro-cognitive symptoms such as nausea, lethargy, and seizures, and, in severe cases, coma and death [3]. Loss of function of any of the urea cycle enzymes causes complete block of ureagenesis, which frequently results in severe hyperammonemia within first few days of life [3, 4]. Approximately 70% of patients with UCD have partial defects in urea cycle enzymes [4] with decreased capacity to produce urea; they can develop hyperammonemia at any time, usually due to infections, invasive medical procedures, fasting or lifestyle changes that result in increased protein catabolism [5-11]. Low protein diet and activation of alternative pathways for disposal of ammonia are standard therapies for urea cycle disorders (UCD) aimed to reduce ammonia production and increase its elimination, respectively [4, 12-14]. Decreased protein intake can be sensed by the body [15], leading to decreased expression of urea cycle genes and enzymes, including the defective one in patients with partial UCD. This can further diminish patient’s capacity to produce urea and increases their risk of hyperammonemia upon sudden increase of protein catabolism. However, this also presents an opportunity for development of new treatments for partial UCD with drugs that increase the expression of urea cycle enzymes despite patients’ low protein diet. Development of such drugs requires detailed understanding of the sensory mechanisms and signaling pathways that regulate expression of the urea cycle genes and enzymes is response to dietary protein intake changes.

Expression of urea cycle genes and abundance of urea cycle enzymes in the liver depend on the rate of protein catabolism, which is determined by the dietary protein intake and the rate of degradation of cellular proteins [15-18]. Glucocorticoids and glucagon act via glucocorticoid receptor and cAMP signaling to activate expression of urea cycle genes [19-26] while the role of insulin in the regulation of urea cycle is unclear [27].

Glucocorticoid receptor regulates expression of the rat *Cps1* gene [28] and cAMP response element binding (CREB) transcription factor regulates rat *Cps1* and human *NAGS* genes [29-31] but transcription factors that regulate expression of each urea cycle gene (Table 1) do not reveal a common regulatory mechanism that might be responsible for coordinated changes in expression of all urea cycle genes in response to changes in dietary protein intake. Therefore, we carried out an integrated analysis of transcriptional and proteomic profiles of the livers from mice fed either high or low protein diet and determined that activity of AMP-dependent protein kinase (AMPK) correlates with expression of urea cycle enzymes in mice. We also examined whether AMPK activates the expression of *OTC* and *CPS1* genes in human primary hepatocytes by exposing them to 5-Aminoimidazole-4-carboxamide ribonucleotide (AICAR).

**Table 1.** Transcription factors that regulate expression of genes that code for urea cycle enzymes.

| Gene | Transcription Initiation | Regulatory Elements |  | References |
| --- | --- | --- | --- | --- |
|  |  | Promoter | Distal Enhancer |  |
| NAGS | Sp1 | CREB | HNF-1, NF-Y | [31, 56] |
| CPS1 | TATA | GR | C/EBP, GR, HNF-3, CREB, P1, P2, P3 | [57-60] |
| OTC | TATA | HNF-4, COUP-TF | C/EBP, HNF-4, COUP-TF | [60-64] |
| ASS | Sp1 | AP2 | CREB | [60] |
| ASL | Sp1 | NF-Y |  | [60] |
| ARG1 | Sp1 | C/EBP, NF-Y, NF-1 | C/EBP, P1<sup>a</sup>, P2<sup>a</sup>, NF-Y | [60, 62, 63] |

<sup>a</sup>The P1 and P2 transcription factors that regulate expression of the ARG1 gene might not be the same as transcription factors designated P1 and P2 that regulate expression of the CPS1 gene.

## Materials and methods

### Time-restricted feeding study

The Institutional Animal Care and Use Committee of the Children’s National Medical Center approved all experimental procedures involving mice.

Groups of eight syngeneic adult male C57BL/6 mice were randomly assigned and fed isocaloric low (20% protein, LP) or high (60% protein, HP) protein diets on a fast/feast cycle consisting of 8 hours fasting, with unlimited access to water, followed by 16 hours *ad libitum* feeding for 11 days. On the 12^th^ day mice were sacrificed immediately before scheduled food reintroduction (T0), and at 30 min. (T30), 60 min. (T60) and 120 min. (T120) after reintroduction of food (Figure S1). Eight mice were sacrificed at each time point and livers were collected and immediately frozen in liquid nitrogen.

### RNA sample preparation and microarray processing

Hierarchical Clustering Explorer (HCE) 3.5 power analysis tool [32] was used to determine that three replicates would be sufficient to allow for detection of significant differences in mRNA levels for 60-80% of probe sets present on each microarray.

RNA was isolated from frozen liver using TRIzol reagent (Invitrogen). The quality of isolated RNA was tested by determining the ratio of absorbance at 260 and 280nm, and visualization of the 28S and 18S ribosomal RNA. Purified RNA was converted to double-stranded cDNA using One-Cycle cDNA Synthesis Kit (Affymetrix) according to manufacturer’s instructions, and cDNA was amplified to ensure greater than 4-fold amplification. The cDNA was transcribed and biotin tagged with the IVT labeling kit (Affymetrix), according to manufacturer’s instructions. The cRNA was purified with the RNAeasy Kit (QIAGEN), and fragmented to approximately 200 bp. To ensure proper fragmentation, the IVT fragmented and un-fragmented samples were visualized on an agarose gel. Following completion of all quality control checks, cRNA was hybridized to GeneChip Mouse Genome 430 2.0 (Affymetrix) and the microarray chips were washed and stained on an Affymetrix Fluidics Station 400 according to manufacturer’s instructions. Fluorescent images were scanned using a Hewlett-Packard G2500A Gene Array Scanner. Hybridization, washing and scanning of chips was carried out within the Center for Genetic Medicine’s Expression Profiling core facility.

### Analysis of microarray data

Affymetrix image data was analyzed using Probe Microarray Suite (MAS) version 5.0 and dCHIP [33] probe set algorithms to highlight genes both algorithms identified as having significant expression level change from the control and to eliminate false positives. Genes identified by each algorithm were imported into Partek software package (Partek Incorporated), normalized by Log_2_, and analyzed by 2-way ANOVA with p-value of p<0.05. Comparisons for the time-restricted feeding study included low protein T0 vs. high protein T0, low protein T30 vs. high protein T30, low protein T60 vs. high protein T60, low protein T120 vs. high protein 120.

ANOVA results were imported into Ingenuity Pathway Analysis (IPA) software using Affymetrix probe set IDs as identifiers and absolute fold-change for observation values. The genes with fold-change cutoffs of less than -1.4 and more than 1.4 were imported into IPA for pathway analysis.

### Validation of gene expression data

RNA was extracted from mouse liver tissues as described previously. Reverse transcription and cDNA synthesis was performed using Oligo-dT 15, random hexamer primers (Invitrogen Life Technologies, Carlsbad, CA) and Superscript III reverse transcriptase (200 U/*μ*g of RNA, Invitrogen Life Technologies) at 42°C for 1 h.

The expression of transcripts with fold changes outside of expression value cutoffs (±1.4-fold) were validated using Applied Biosystems 384 custom gene card array. 64 genes (Table S1) were assayed by quantitative PCR using 7900HT Fast Real-Time PCR System (Applied Biosystems, Inc.) and included myosin heavy chain 9 as the normalization control that does not change due to diet and 18S RNA as the manufacturer’s control. The ΔΔCt method was used to calculate the difference in expression levels for each gene [34].

### Liver proteome profiling by mass spectrometry

Protein lysates were prepared from frozen mouse liver tissue by homogenization of tissue in 50mM Tris-acetate buffer, pH 7.5 containing 250mM sucrose, 1mM EDTA, 1% Nonident P40 (NP40), and Complete Mini Protease Inhibitor Cocktail (Roche). Aliquots of 50 *μ*g of total liver proteins from wild-type mice labeled with ^13^C_6_-lysine [35] were added to 50 *μ*g of total liver proteins from each experimental mouse. Samples containing stable isotope labeled and unlabeled protein mixture were resolved by SDS-PAGE, stained with Bio-Safe Coomassie (Bio-Rad, Hercules, CA), and 1mm wide sections were excised from each gel lane. Proteins in excised gel slices were digested with trypsin as previously described [36]. Concentrated peptides from each band were injected via an autosampler (6 *μ*l) and loaded onto a Symmetry C18 trap column (5μm, 300 μm i.d. × 23 mm, Waters) for 10 min. at a flow rate of 10 μL/min, 0.1% formic acid. The samples were subsequently separated by a C18 reverse-phase column (3 μm, 200A, 100 μm × 15 cm, Magic C18, Michrom Bioresources) at a flow rate of 300 nl/min. using an Eksigent nano-hplc system (Dublin, CA). The mobile phases consisted of water with 0.1% formic acid (A) and 90% acetonitrile (B). A 65 min. linear gradient from 5 to 60% B was employed. Eluted peptides were introduced into the mass spectrometer via Michrom Bioresources CaptiveSpray. The spray voltage was set at 1.4 kV and the heated capillary at 200°C. The LTQ-Orbitrap-XL (ThermoFisherScientific) was operated in data-dependent mode with dynamic exclusion in which one cycle of experiments consisted of a full-MS in the Orbitrap (300-2000 m/z) survey scan in profile mode, resolution 30,000, and five subsequent MS/MS scans in the LTQ of the most intense peaks in centroid mode using collision-induced dissociation with the collision gas (helium) and normalized collision energy value set at 35%.

### Database search and SILAM ratio measurement

For protein identification and quantification we used Integrated Proteomics Pipeline (IP2) version 1.01 software developed by Integrated Proteomics Applications, Inc. (https://www.integratedproteomics.com/). LC-MS/MS raw data were uploaded into IP2 software. Files from each lane were searched against the forward and reverse Uniprot mouse database (UniProt release 15.4, June 2009) for tryptic peptides allowing one missed cleavage, and possible modification of oxidized methionine (15.99492 Da) and heavy lysine (6.0204 Da). IP2 uses the Sequest 2010 (06 10 13 1836) search engine. Mass tolerance was set at ±30 ppm for MS and ±1.5 Da for MS/MS. Data were filtered based on a 3% false discovery rate. All the bands from each lane were summed in the analysis. Census software version 1.77, built into the IP2 platform, was used to determine the ratios of unlabeled and labeled peptide pairs using an extracted chromatogram approach. The distribution of ratios was plotted and correction factors applied to adjust for error in sample mixing. Data were checked for validity by using regression correlation better than 0.98 for each peptide pair.

### Validation of protein profiling data

Mouse liver tissue was homogenized in an extraction buffer containing 50mM Tris-acetate pH 7.5, 250 mM sucrose, 1mM EDTA, 1% Nonidet P40 and anti-protease mixture (Roche), and diluted to a protein concentration of 20 mg/ml. NAGS was probed in 20 *μ*g of total liver proteins, which were resolved by SDS-PAGE and transferred to a nitrocellulose filter. Filters were blocked with blocking buffer (Pierce) containing 0.5% Surfact-Amps 20 (Pierce). The filter was probed with polyclonal antibody raised against recombinant mouse NAGS at 1:5,000 dilution for 1 hr. at room temperature and washed with Tris buffered saline containing 0.005% Tween-20 (TBST). The filter was then incubated for 1 hr. at room temperature with donkey horseradish peroxidase (HPRT). NAGS bands were visualized using SuperSignal West Pico kit (Pierce) according to the manufacturer’s instructions. CPS1 was probed in 150 ng of total liver protein using rabbit anti-CPS1 primary antibody (AbCam) at 1:5000 dilution and HPRT-conjugated donkey anti-rabbit secondary antibody (Pierce). OTC was probed in 250 ng of total liver protein using rabbit primary antibody raised against recombinant OTC at 1:5000 dilution and HPRT-conjugated donkey anti-rabbit secondary antibody (Pierce). ASS was probed in 250 ng of total liver protein using goat anti-ASS primary antibody (Santa Cruz Inc.) at 1:500 dilution and HPRT-conjugated donkey anti-goat secondary antibody (Promega). ASL was probed in 40 *μ*g of total liver protein using goat anti-ASL primary antibody (Santa Cruz Inc.) at 1:500 dilution and HPRT-conjugated donkey anti-goat secondary antibody (Promega). Vinculin was probed with mouse anti-vinculin primary antibody (Sigma) at 1:1000 dilution and HPRT-conjugated goat anti-mouse secondary antibody (Bio-Rad) at 1:2000 dilution. CPS1, OTC, ASS, ASL and vinculin were visualized with the ECL Western Blotting Substrate (Pierce) according to the manufacturer’s instructions.

### Measurement of protein phosphorylation

Proteins were extracted from liver tissue using cell lysis buffer (Cell Signaling Technology) according to manufacturer’s instructions with the modification for lysis of tissue samples. Briefly, 100mg of liver tissue was incubated with 2x lysis buffer and sonicated using 3 pulses of 15 sec. at 30% output. Tissue suspension was centrifuged and supernatent was collected for analysis. 30 *μ*g of whole liver lysate was separated under denaturing conditions by 4-20% SDS-Page gel electrophoresis and transferred to PVDF membranes. Membranes were blocked in 50% Blocking Buffer (Odyssey) and labeled with primary rabbit antibodies to phosphorylated and non-phosphorylated AMPKa, mTOR, 4E-BP1 and *β*-actin proteins (Cell Signaling Technologies) at a 1:1000 dilution. The membranes were washed three times for 10 min, each with TBS containing 0.1% Tween 20, followed by incubation with secondary antibody at a 1:10,000 dilution. Image analyses were performed using an Odyssey Imager (LiCor, Lincoln, NE).

### Primary hepatocyte culture

The Liver Tissue Cell Distribution System (LTCDS) is a National Institutes of Health (NIH)-funded collaborative network to provide human liver tissue from regional centers for distribution to scientific investigators. The regional centers have active liver transplant programs with human subjects’ approval to provide portions of the resected pathologic liver for which the transplant is performed. All human tissues were collected with informed consent following ethical and institutional guidelines. Tissue dissociation and subsequent hepatocyte isolation procedures were developed by Seglen with modifications [37]. Hepatocytes were allowed to adhere for 2 hours with Hepatocyte Maintenance Media (Lonza, Walkersville, MD) containing 10% FBS and penicillin and streptomycin. This was followed by maintenance culture for 1 days in serum-free Hepatocyte Maintenance Media (Lonza, Walkersville, MD). Hepatocytes were treated with 5 μM AICAR with or without 40 μM Compound C (Sigma-Alrich, St. Louis, MO) for 24 hours. Cell viability was determined using PrestoBlue Cell Viability Reagent (Life Technologies, Grand Island, NY) according to the manufacturer’s instructions. Cell pellets and supernatants were stored at -80°C until analysis.

### Quantitative RT-PCR

RNA was extracted from cell pellets using RNeasy Minikit (Qiagen) according to the manufacturer’s instructions. One microgram of RNA was reverse transcribed to cDNA using a modified MMLV-reverse transcriptase with RNase H^+^ activity (iScript, Bio-Rad, Hercules, CA) according to the manufacturer’s instructions. Real-time quantitative PCR reactions were carried out in 50 *μ*L using TaqMan systems (Applied Biosciences, Carlsbad, CA) using ABI 7500 Fast Real Time PCR System (Applied Biosystems, Foster City, CA).

## Results

### Physiological effects of time-restricted feeding on mice

The body weight of mice fed either high (60%) or low (20%) protein (HP and LP, respectively) diet did not differ at the beginning and at the end of the 12 days of time-restricted feeding (Figure 1A). Mice fed HP gained weight during experiment while there was a trend towards higher weight at the end of experiment in mice fed LP (Figure 1A). Food intake was lower in mice fed the high protein diet (Figure 1B), most likely due to the prolonged satiety following consumption of high protein food [38, 39]. Mice fed high protein diet drank more water, presumably to allow for increased urea excretion through urination (Figure 1C). While urea excretion was not measured, the high protein diet would result in increased urea production [40].

**Figure 1.**
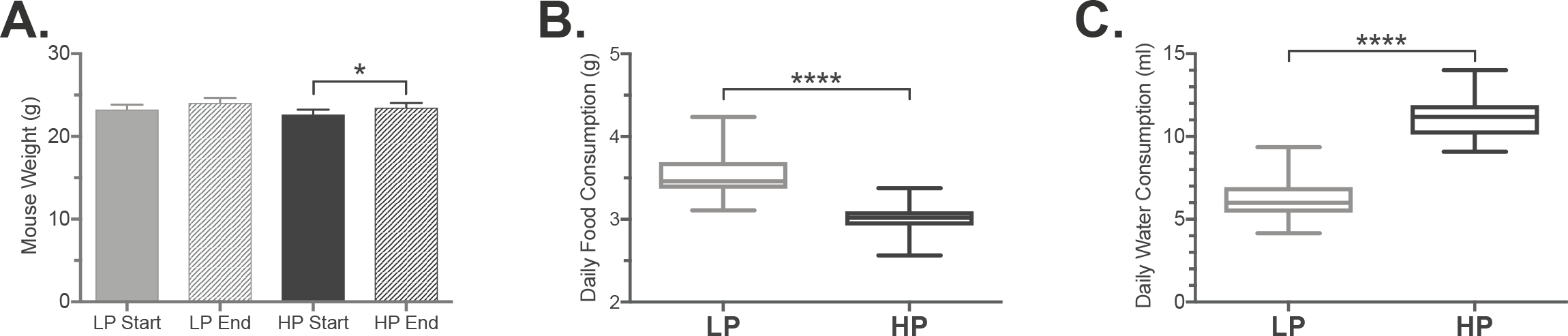
Physiological effects of different protein intake. Mice were fed either LP (light gray) or HP (dark gray) diets on a time-restricted schedule. **A.** The average weight of animals at the beginning (solid) and the end (hatched) of experiment. **B.** Average daily food consumption of mice fed either LP or HP. **C.** Average daily water consumption of mice fed either LP or HP. **** – p<000.1.

### Proteomic and transcriptional profiling

Livers were harvested from mice fed either HP or LP diet on a time-restricted feeding schedule of 6 hr. fasting and 18 hr. feeding for 11 days. On the 12^th^ day, livers were collected immediately prior to reintroduction of food, and 30, 60, and 120 minutes after reintroduction of food (T0, T30, T60 and T120 time points, respectively); expression microarrays were used to assess mRNA levels at each time point and spike-in differential proteome profiling was used to detect changes in protein abundance in mice fed either LP or HP diets.

Differences in expression levels of urea cycle mRNA and proteins were assessed first. Expression levels of *Cps1*, *Otc*, *Ass*, and *Asl* mRNA were higher in mice fed HP diet, while the abundance of the *Nags* and *Arg1* mRNA did not change in response to diet (Figures 2A and 2B). The Cps1, Otc, Ass, Asl and Arg1 proteins were more abundant in the livers of mice fed HP diet (Figures 2C-2E), similar to changes in their abundance and enzymatic activity in response to HP diet previously observed [18, 41]. The Nags protein was assayed only by immunoblotting because it could not be detected by mass spectroscopy due to its very low abundance. Unlike other urea cycle proteins, the Nags was more abundant in the livers of mice fed LP diet than in the livers of mice fed HP diet (Figures 2D and 2E).

**Figure 2.**
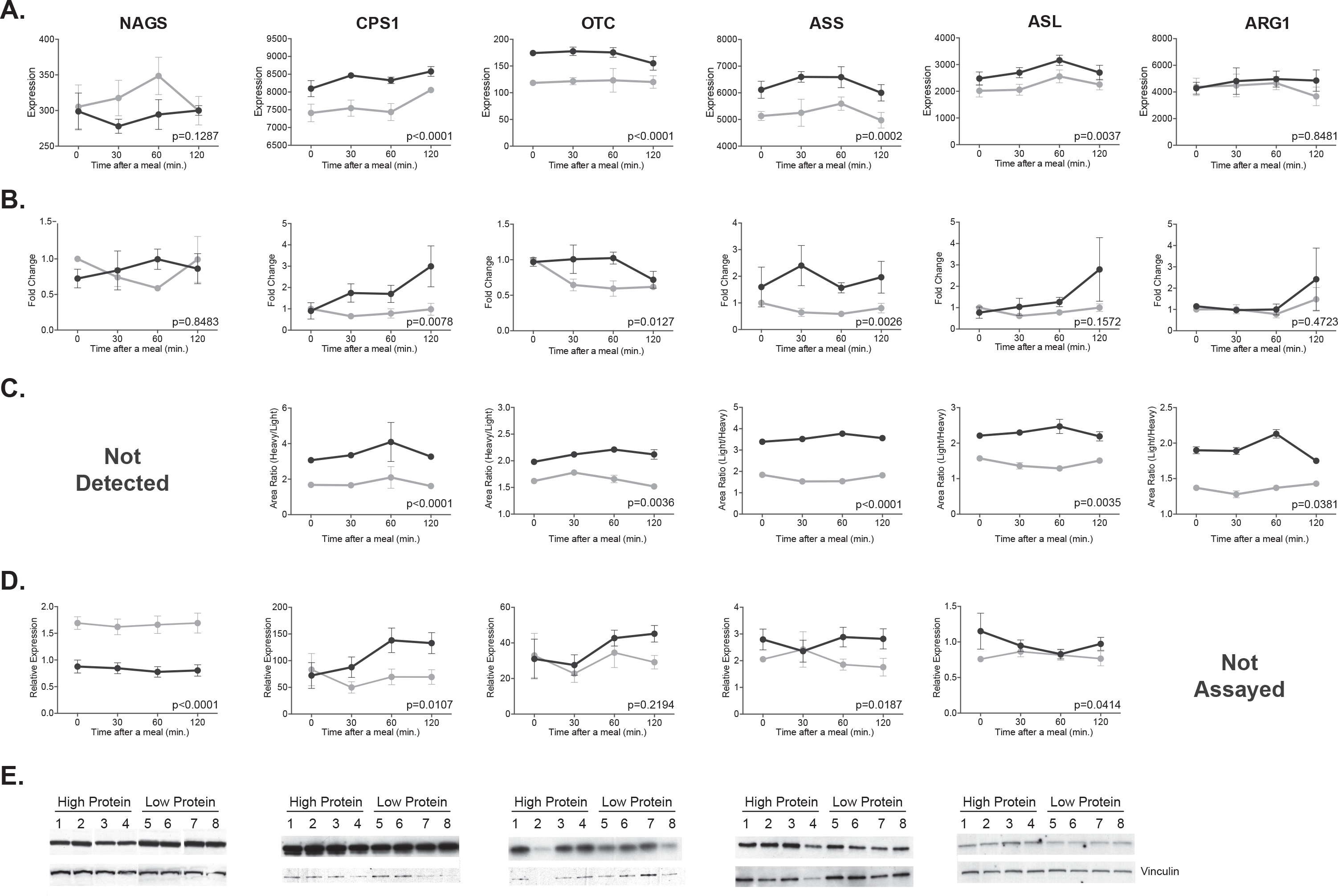
Expression of urea cycle genes and enzymes in mice fed either HP (dark gray) or LP (light gray) diets. The mRNA abundance was measured using Affymetrix microarrays (**A**) and validated with quantitative RT-PCR in a different cohort of mice (**B**). The abundance of urea cycle enzymes was measured using spike-in mass spectrometry (**C**) and validated with immunoblotting in a different cohort of mice (**D**and **E**). Abundance of mRNA and proteins was measured at fasting and 30, 60 and 120 min. after introduction of food. Each data point is a mean and associated SEM of 4 measurements.

Of the 45101 probe sets used for analysis, 505, 604, 740 and 447 probe sets indicated expression fold change ±1.4 or more and p<0.05 in mice fed HP and LP at T0, T30, T60 and T120, respectively (Figure S2). Proteomic analysis was initially done on a single mouse fed each diet at each time point. Because none of the detected proteins changed abundance over time we focused on analyzing differences in protein abundance at T0. Of the 1073 proteins that were identified and quantified in mice fed HP and LP at T0, 335 had fold change ±1.2 or more and p<0.05 (Figure S3). When lists of genes that were differentially expressed were analyzed using Integrated Pathway Analysis software package, there were no obvious signaling or metabolic pathways that responded to dietary protein intake. We then looked simultaneously at ±1.5-fold changes in mRNA and protein abundance in mice fed HP and LP at T0 (Figure 3). Abundance of mRNA and protein for aldehyde dehydrogenase X (AdhX) and glycine dehydrogenase (Gldh) was higher in mice fed HP, while 11 mRNA and proteins were down regulated in these mice (Figure 3 and Table S2). 70 proteins were at least 1.5-fold more abundant in the livers of mice fed HP while 24 proteins were at least 1.5-fold less abundant, while the abundance of their mRNA did not change (Figure 3 and Table S2). Many of the genes and proteins that were up or down-regulated in mice fed HP participate in metabolism of amino acids, carbohydrates, fats and steroids (Figure 4). Defects in many proteins that are up-regulated in mice fed HP cause inborn errors of metabolism. Absence or reduction in activity of *PCCA* and *PCCB* cause propionic academia [42]; maple syrup urine disease is caused by defects in *BCKDHA* and *BCKDHB* genes and proteins [43], metymalonyl academia is caused by defects in the *MUT* gene and protein [44].

Although mice fed HP and LP had the same fat intake, the acetyl CoA carboxylase *α* (Acc*α*), fatty acid synthase (Fasn) and 3-hydroxy-3-methyl-glutaryl-coenzyme A reductase (Hmgcr), the key enzymes of the fatty acid and steroid biosynthesis [45-47] were down regulated in mice fed HP (Figure 4). This prompted us to validate mRNA expression changes of these three genes and to examine expression levels of *Ampkα* mRNA because it regulates activity and expression of the three enzymes [45, 48, 49].

Fluorescence intensity data of probe sets that correspond to *Ampkα*, *Accα*, *Fasn* and *Hmgcr* were extracted from the data generated by PLIER, and the difference in abundance of these four mRNAs was measured by quantitative PCR in a separate cohort of mice fed HP and LP. Profiling and qPCR data indicated that expression of the *Ampkα* mRNA did not change in response to diet (Figure 5A). The gene expression array indicated lower abundance of the *Ampkα* mRNA in the livers of mice fed HP diet while validation of expression data by quantitative RT-PCR revealed only a trend towards lower expression of the *Accα* mRNA in the livers of mice fed HP diet (Figure 5B). The abundance of the *Fasn* and *Hmgcr* mRNA was lower in the livers of mice fed HP (Figures 5C and 5D).

**Figure 3.**
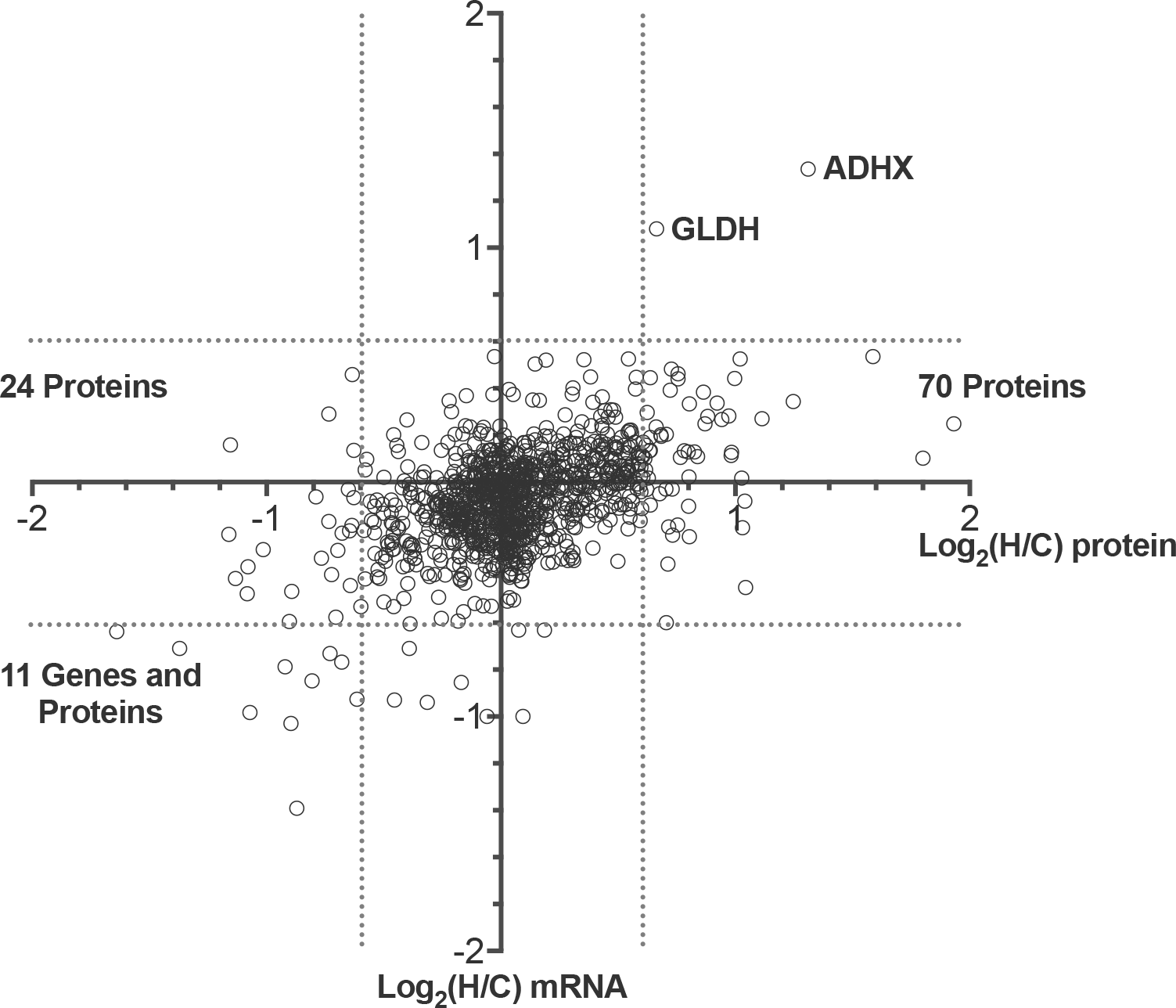
Correlation between changes in abundance of mRNA and proteins in livers of mice fed either HP or LP diets. ADH – aldehyde dehydrogenase X, GLDH – glycine dehydrogenase. Hatched lines indicate ±Log_2_1.5.

**Figure 4.**
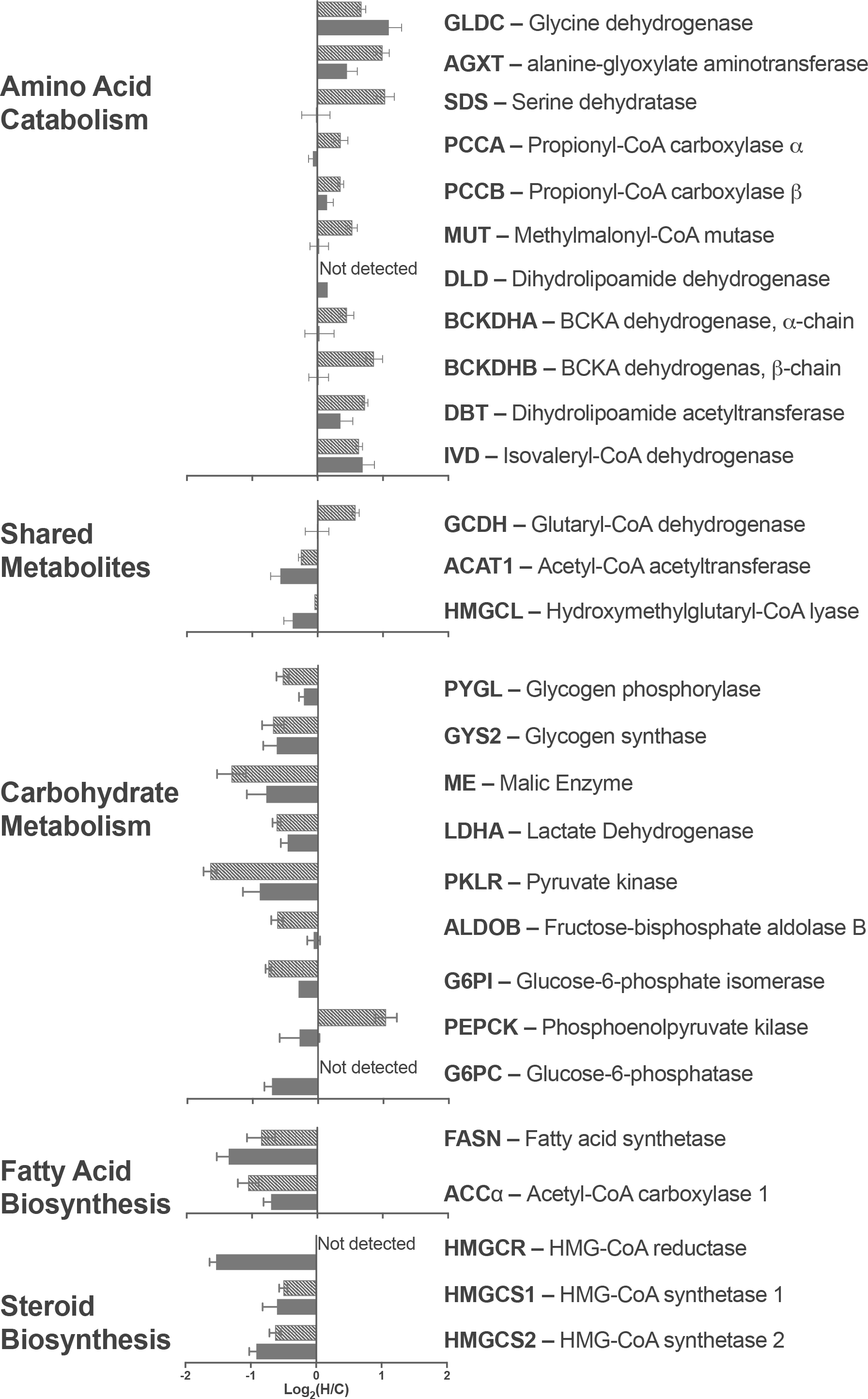
Expression changes of mRNA and enzymes in amino acid catabolism, carbohydrate metabolism, and fatty acid and steroid biosynthesis in mice fed either HP or LP diets. Solid – mRNA expression. Hatched – Protein expression.

**Figure 5.**
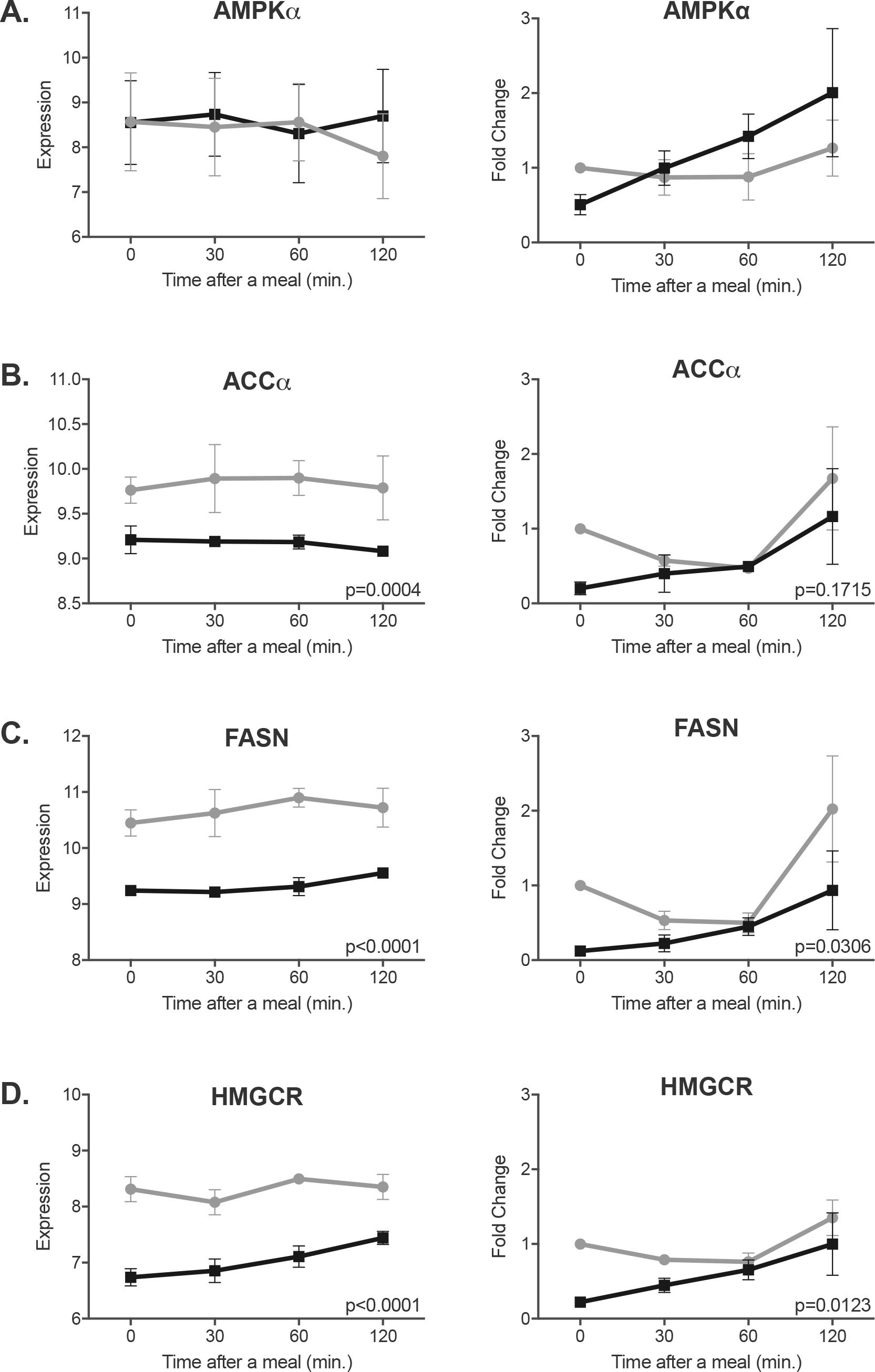
Expression changes of AMPKα (**A**), ACCα (**B**), FASN (**C**) and HMGCR (**D**) mRNA in the livers of mice fed either HP (dark grey) or LP (light gray) diets. Abundance of mRNA was measured at fasting and 30, 60 and 120 min. after introduction of food. Graphs on the left show expression measurements using oligonucleotide microarrays. Graphs on the right show expression measurements using quantitative RT-PCR in a different cohort of mice. Each data point is a mean and associated SEM of 4 measurements.

### Activated AMPK signaling in the livers of mice fed HP diet

Because AMPK signaling regulates expression and enzymatic activity of Acc*α* Fasn and Hmgcr [45, 48, 49] we examined whether phosphorylation levels of Ampk*α* and its downstream targets, mammalian target of rapamycin (mTor) and eukaryotic translation initiation factor 4E binding protein 1 (4E-BP1), differ at fasting and at different times after a meal in the livers of mice fed HP and LP. Liver proteins from mice fed HP and LP were resolved using SDS-PAGE and probed with antibodies that detect phospho-Thr^172^ in Ampk phospho-Thr^2446^ in mTor, and phospho-Thr^37^ and Thr^46^ (Thr^37/46^) in the 4E-BP1, followed by immunoblotting with antibodies that detect the total amount of the three proteins. Beta-actin was used as a loading control (Figure S4). Phosphorylation levels of each protein were calculated from the ratio of band intensities of the phosphorylated and total protein and normalized to the *β*-actin levels. There was a trend (p=0.097) toward different pattern of Thr^172^ phosphorylation levels of the Ampk*α*in mice fed HP and LP. The fasting levels of Thr^172^ phosphorylation levels were lower in mice fed HP and they tended to increase thereafter, while phosphorylation of Thr^172^ decreased during and after a LP meal (Figure 6A). Patterns of phosphorylation changes of mTor and 4E-BP1 were similar. Phosphorylation of levels Thr^2446^ and Thr^37/46^ in the mTor and 4E-BP1, respectively, did not change during and after an LP meal. However, in mice fed HP diet, phosphorylation of the mTor Thr^2446^ and 4E-BP1 Thr^37/46^ at fasting was lower than in mice fed LP diet; 30 and 60 min. after introduction of the HP meal, phosphorylation levels of the mTOR Thr^2446^ and 4E-BP1 Thr^37/46^ increased and then 120 min. after introduction of the HP meal, decreased to levels lower than in mice fed LP diet (Figures 6B and 6C). These differences in the patterns of phosphorylation levels of the Ampk*α* mTor and 4E-BP1 may be responsible for changes in the abundance of mRNA and proteins of the urea cycle enzymes and adaptation to HP diet.

**Figure 6.**
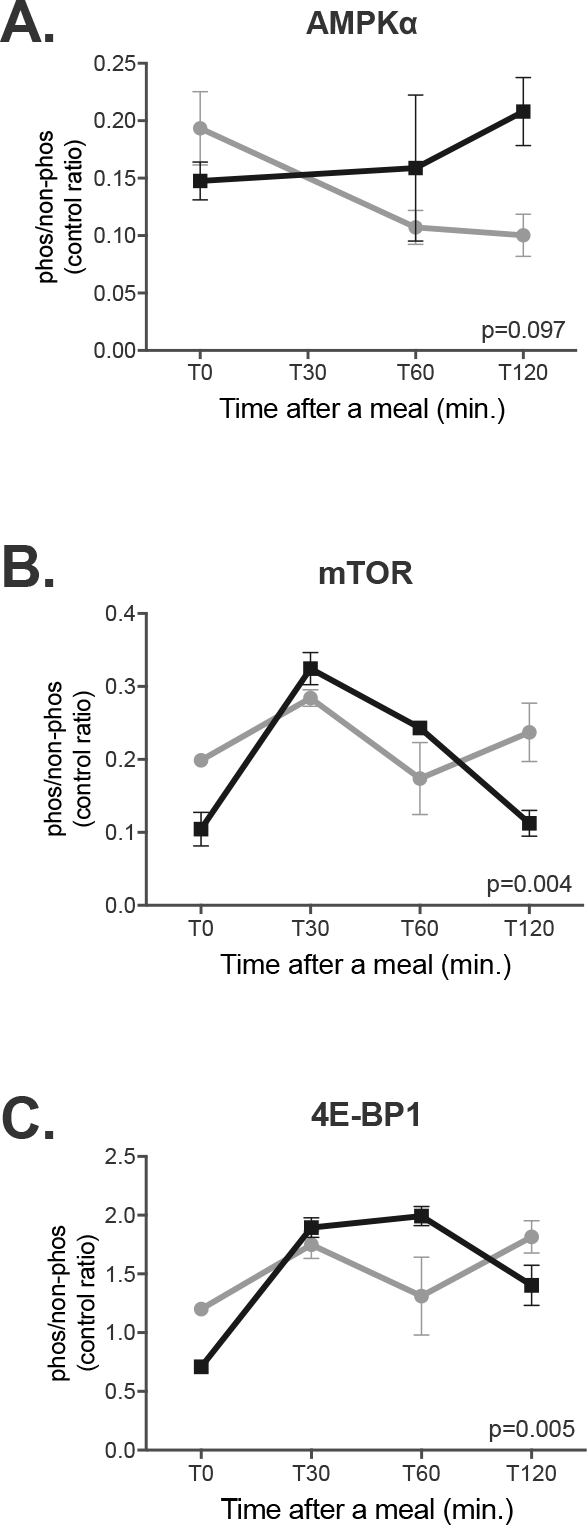
Changes in phosphorylation of AMPK (**A**) and its targets mTOR (**B**) and 4E-BP1 (**C**) in the livers of mice fed either HP or LP diets. Phosphorylation was measured at fasting and 30, 60 and 120 min. after introduction of food. The ratio of phosphorylated and non-phosphorylated protein was normalized to [⊠-actin abundance. Each data point is a mean and associated SEM of either 4 (AMPK) or 3 (mTOR and 4E-BP1) measurements.

### Activation of AMPK*α* increases expression of the *CPS1* and *OTC* mRNA

Inverse correlation of expression changes of the urea cycle enzymes and the key enzymes in the fatty acid and steroid synthesis raised the possibility that AMPK signaling may regulate expression of urea cycle enzymes in response to changes in the dietary protein intake. We used human primary hepatocytes to test whether activation of AMPK signaling results in increased expression of the CPS1 and OTC mRNA. Primary human hepatocytes were treated either with AICAR, which activates AMPK, or with AICAR and compound C, a reversible inhibitor of AMPK, or with PBS as a control, followed by measurements of cell viability and the *CPS1* and *OTC* mRNA levels. These treatments did not affect viability of the primary human hepatocytes (Figure 7A). Expression levels of *CPS1* and *OTC* mRNA increased 1.8- and 2.8-fold, respectively, in primary human hepatocytes treated with AICAR (Figures 7B and 7C). Expression of the *CPS1* mRNA decreased two-fold in cells treated with AICAR and compound C, while the expression levels of the *OTC* mRNA remained unchanged (Figures 7B and 7C). These data strongly suggest that AMPK signaling regulates expression of at least two urea cycle enzymes.

**Figure 7.**
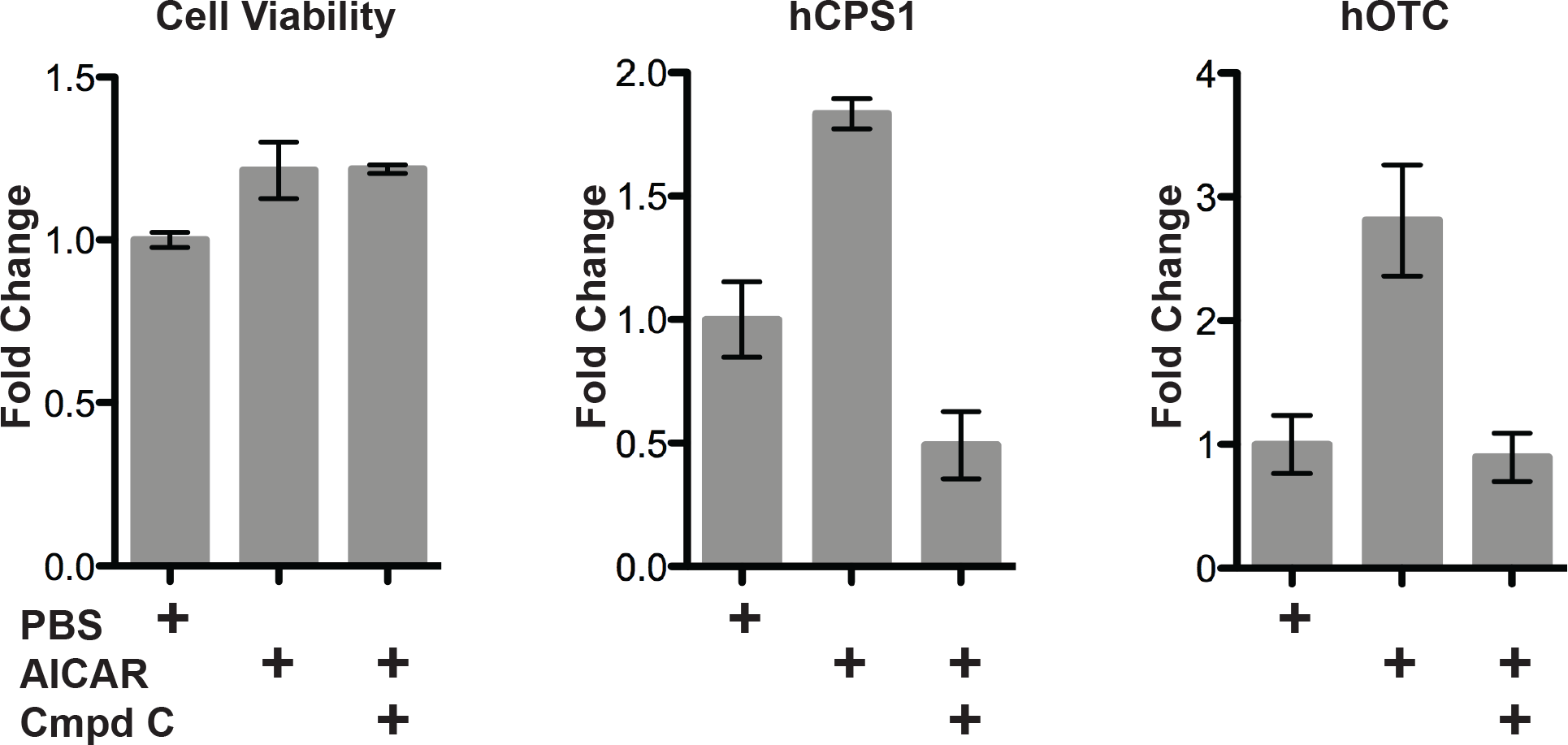
Cell viability (**A**) and expression of *CPS1* (**B**) and *OTC* (**C**) mRNA in primary human hepatocytes treated with either AICAR or AICAR and compound C. Each data point is a mean and associated SEM of 3 measurements.

## Discussion

The goal of this study was to identify signaling pathways that regulate expression of urea cycle genes in response to changes in the dietary protein intake. We used integrated transcriptional and proteomic profiling to show that activation of the AMPK signaling correlates with increased expression of urea cycle genes in mice fed HP diet. A direct role of AMPK in increased expression of at least two urea cycle genes, *CPS1* and *OTC*, was confirmed in human primary hepatocytes treated with AMPK activator AICAR. The decreased abundance of the Nags protein in response to HP diet was unexpected and may indicate a different regulatory mechanism for expression of the mouse *Nags* gene. Enzymatic activity of mammalian recombinant NAGS doubles upon binding of L-arginine [50, 51] and intake of this amino acid would be higher in mice fed HP diet. Therefore, it is possible that higher intake of L-arginine in mice fed HP diet activated Nags and resulted in higher production of NAG for activation of CPS1 despite lower abundance of the Nags enzyme. While this mechanism may be responsible for short-term regulation of urea production it may not explain adaptive changes in expression of urea cycle genes in response to dietary protein intake.

Integration of transcription and protein profiles was needed because neither approach alone pointed to a signaling pathway that could be responsible for changes in expression of urea cycle genes and abundance of urea cycle enzymes in response to increased dietary protein intake. This is not surprising since changes in dietary protein intake are more likely to trigger changes in activity and phosphorylation status of various signaling proteins than their abundance or expression of their mRNA. Combined mRNA and protein expression analysis identified coordinated decrease in the abundance of the key enzymes in the fatty acid and steroid biosynthesis in mice fed HP diet, a process that is regulated by AMPK signaling [45, 48, 49, 52]. This suggested that AMPK might also regulate expression of the urea cycle genes and enzymes in response to dietary protein intake while increased expression of the *CPS1* and *OTC* mRNA in primary human hepatocytes treated with AICAR, and activator of AMPK, supports its role in regulation of expression of at least two urea cycle genes. Our results are also consistent with the possibility that regulation of urea cycle genes and enzymes was in response to low carbohydrate instead of high protein intake because the fraction of starch was varied in the diets of experimental animals depending on the protein content.

The patterns of the AMPK Thr^172^ phosphorylation differed in mice fed either HP or LP diet. Because Thr^172^ can be phosphorylated by the liver kinase B1 (LKB1), which can be activated by the protein kinase A (PKA) and cAMP [53], and cAMP and PKA also activate expression of urea cycle genes [19, 20, 22, 24-26, 41, 54] we propose that AMPK could be downstream of cAMP and PKA in the signaling cascade that regulates expression of urea cycle genes in response to diet.

A role for AMPK in the regulation of urea cycle enzymes opens up new prospects for treatment of patients with partial defects of UCD, who have diminished capacity to produce urea and represent about 70% of all UCD patients [4, 55]. Protein restricted diet is a mainstay treatment for patients with UCD because it lowers production of ammonia in their bodies. Because our bodies can sense the amount of dietary protein intake and adjust expression of urea cycle genes and enzymes accordingly, the low protein diet can put these patients at increased risk of hyperammonemia due to further decrease of their already low capacity to produce urea. A drug treatment targeted at increasing expression of urea cycle genes and enzymes, including the partially defective one, despite a low protein diet, could benefit patients with partial UCD by increasing their capacity to produce urea and decreasing the risk of hyperammonemia. Because of its central role in cellular energy metabolism, AMPK has been implicated in diseases such as diabetes, cancer and obesity, and has been a target for drug development [55] that also might benefit patients with UCD.

## Acknowledgments

Sree Rayavarapu performed these studies at the Children’s National Medical Center as part of their doctoral studies. This work was supported by public health service grants K01DK076846 and R01DK064913 from the National Institute of Diabetes Digestive and Kidney Diseases, National Institutes of Health, Department of Health and Human Services.

## Disclaimer

The views expressed in this manuscript are those of the authors and do not reflect the official policy of the FDA. No official support or endorsement by the FDA is intended or should be inferred.

## Author contributions

SKH performed time-restricted feeding study, microarray analysis of mRNA expression, validation of the expression data, and immunoblotting of urea cycle enzymes and wrote the manuscript. PJM measured CPS1 and OTC mRNA expression in primary human hepatocytes and phosphorylation levels of AMPK*α*, mTOR and 4E-BP1 in mouse livers. NH prepared liver proteins for proteomic analysis. HDM carried out quantitation of the NAGS protein using immunoblotting. SR and KN generated ^15^N-labeled mice for the spike-in differential proteomic profiling. YH and KB carried out quantitation of liver protein using mass spectroscopy, database search and SILAM ratio measurement. MT participated in data interpretation and critically reviewed the manuscript. LC conceived of the study, carried out integrated analysis of the transcription and proteomic profiles and edited the manuscript

## Supporting information

**S1 Fig.** Experimental design for identifying signaling pathways that regulate expression of urea cycle genes.

**S2 Fig.** Scatter plot of statistical significance vs. fold change of mRNA expression in the livers of mice fed either HP or LP diets.

**S3 Fig.** Scatter plot of statistical significance vs. fold change of protein expression in the livers of mice fed either HP or LP.

**S4 Fig.** Quantification of phosphorylated AMPK*α* subunit, mTOR and 4E-BP1 using immunoblotting.

**S1 Table.** List of genes that were included in the Applied Biosystems 384 custom gene card array.

**S2 Table.** List of genes/probe sets that had -1.4≥fold change≥1.4 and p<0.05 at T0, T30, T60 and T120.

**S3 Table.** List of proteins that had -1.2≥fold change≥1.2 and p<0.05 at T0.

**S4 Table.** List of genes and proteins that were included in the integrated analysis of transcriptional and protein profiles.

